# Antimicrobial peptide databases and prediction tools: Toward a standard evaluation framework

**DOI:** 10.64898/2026.05.19.726290

**Authors:** Alejandro Cisterna-García, Alvaro M. González-López, Aspasia Vozi, María A. Esteban, Adrian Egli, Catherine Jutzeler, Jose Palma, Alvaro Sánchez-Ferrer, Juan A. Botía

## Abstract

Antimicrobial resistance (AMR) has a profound impact on animal and human health and is associated with substantial morbidity, mortality and public health costs. There is a clear need to develop novel, effective antibiotic agents, which can overcome the current AMR crisis. Antimicrobial peptides (AMPs) may offer such a solution and have attracted growing attention for their potential to combat AMR. In parallel, the growing availability of peptide sequences in public databases has stimulated the development of numerous machine learning and deep learning tools to predict antimicrobial activity computationally. However, it remains unclear how reliably these tools can be compared, as existing studies often rely on heterogeneous datasets and inconsistent evaluation protocols that may lead to data leakage and inflated performance estimates. This raises a central question: what evaluation criteria and benchmark resources are needed to enable fair, reproducible, and biologically meaningful assessment of AMP prediction tools? We address this question by focusing specifically on antibacterial peptides (ABPs). We first provide an overview of AMP databases relevant to antibacterial activity and compare their content, redundancy, and experimental metadata. We then critically assess existing computational tools for ABP prediction, highlighting key limitations related to dataset construction, affinity to certain sequences, data leakage, and inconsistent performance reporting. Based on these limitations, we propose a reference evaluation framework designed to improve comparability, reproducibility, and practical utility in ABP prediction. Finally, we provide targeted recommendations for AMP databases and future tool development to support more robust progress in the computational discovery of ABPs.

## Introduction

The number of deaths caused by complications derived from infections was around 13.7 millions in 2019 ^1^. Of these, 4.7 million were associated with pathogens carrying antimicrobial resistance (AMR) pathogens, often referred to as ‘superbug’ ^2^. In response to this growing threat, the World Health Organization (WHO) included AMR in its list of urgent global health challenges in January 2020, highlighting the critical need for new treatments ^3^ among other counter measurements. Furthermore, WHO also published a list with their bacterial priority pathogens ^4^. The WHO report states that more efforts and investments in novel antimicrobials are needed to address these most impactful pathogens, which include high-burden AMR bacteria such as Enterobacterales, particularly carbapenemase-producing Enterobacterales (CPE); non-fermenting Gram-negative bacteria, including Pseudomonas aeruginosa and Acinetobacter baumannii; Salmonella and Shigella spp.; Neisseria gonorrhoeae; and Staphylococcus aureus. Antimicrobial peptides (AMPs) emerge as a potential source of new treatments against these highly resistant bacteria ^5–8^.

AMPs are short amino acid sequences that are generally considered to be less than 50 amino acids in length ^9^, and are found in a wide variety of organisms, ranging from single-celled prokaryotes to multicellular eukaryotes, including insects, amphibians, plants, and humans ^10–12^. AMPs have been routinely collected and classified in various databases, such as the Antimicrobial Peptide Database (APD) ^13^, dbAMP ^14^, and the Antimicrobial Peptide Data Repository (DRAMP) ^15^, based on their biological properties, including their antibacterial, antifungal, antiviral, and antiparasitic activities ^16–18^. The classical process of AMP discovery is complex, multi-step, time-consuming, and resource-intensive. It usually begins with the identification of organisms, such as bacteria, fungi, plants, or animals, that produce bioactive peptides ^19–21^. Candidate peptides are then isolated from biological extracts or predicted from genomic, transcriptomic, or proteomic data, followed by purification and structural characterization using techniques such as chromatography and mass spectrometry ^22,23^. Once identified, these peptides must be synthesized or extracted in sufficient quantities and experimentally validated through in vitro assays against target pathogens ^24^. These assays conceptually assess pathogen growth in presence of increasing concentration of the AMP. Promising candidates undergo further evaluation for stability, toxicity, selectivity, and mechanism of action, often requiring iterative rounds of optimization through sequence modification and structure–activity relationship studies ^25–27^. As a result, this traditional pipeline demands substantial experimental effort, specialized infrastructure, and long development timelines.

As an alternative to this tedious process, a series of computational tools have been developed that employ machine learning (ML) and deep learning (DL) approaches, leveraging the pattern-recognition capabilities of algorithms to identify patterns useful for distinguishing AMPs from non-AMPs ^28–53^. Typically, these predictive models are trained using annotated AMP sequences from AMP-specific databases and non-AMP sequences retrieved from general protein repositories, such as UniProt. The model’s performance is then typically evaluated using independent datasets of peptide sequences from other AMP databases. However, despite the proliferation of prediction tools based on ML and DL, the lack of a common dataset which can be used as a datasets across different tools makes direct comparison difficult and complicates the selection of the most appropriate tool.

Addressing existing research gaps in AMP discovery, this review focuses exclusively on antibacterial peptides (ABPs), prioritized for their clinical relevance and relative structural homogeneity. We introduce a groundbreaking benchmark that utilizes a comprehensive, integrated dataset of ABP sequences compiled from the current database ecosystem (**Figure 1**). By systematically evaluating diverse AMP predictive tools against this unified and curated set, we draw attention to the critical limitations of existing models and propose a roadmap to refine predictive accuracy in this field.

**Figure 1.**
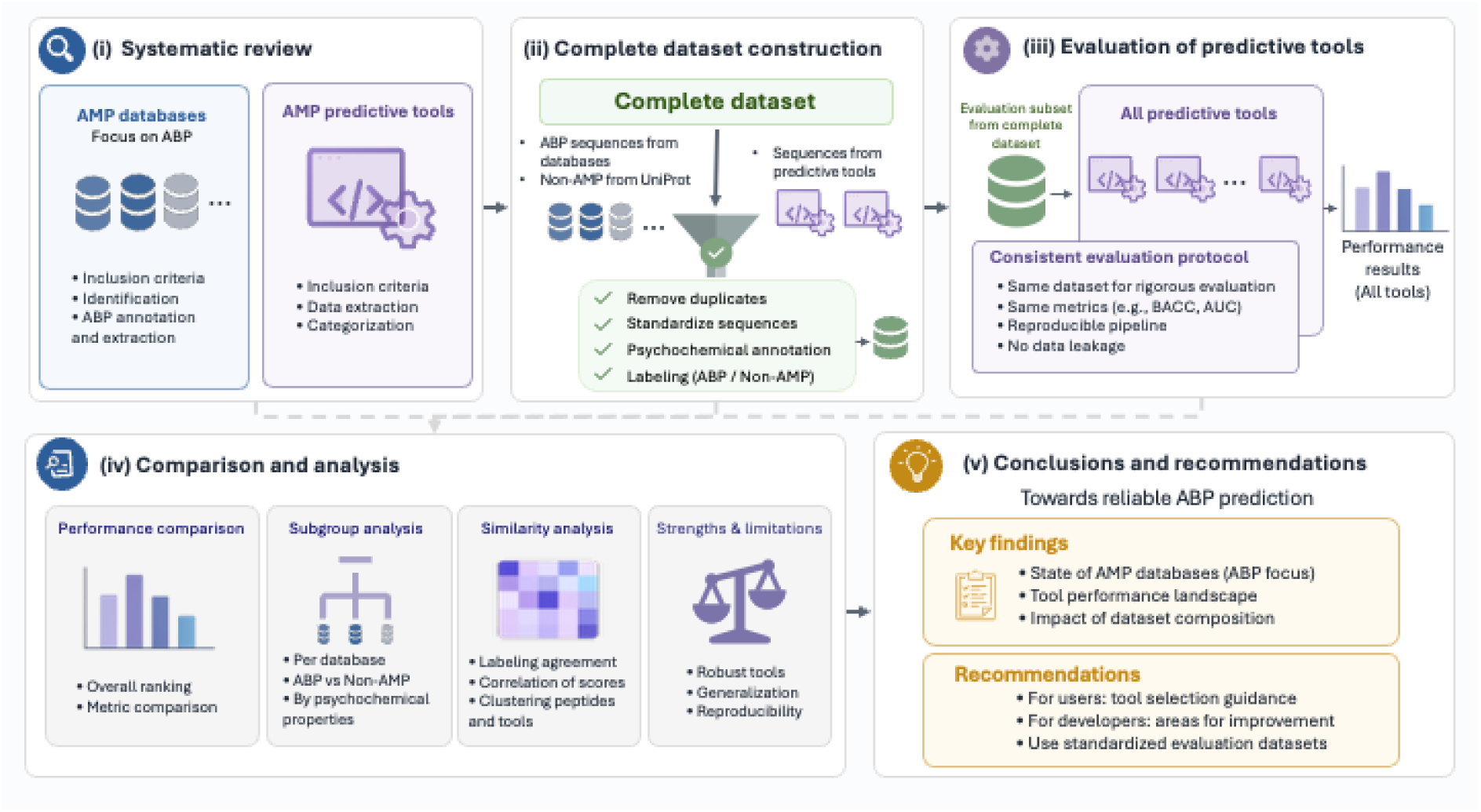
Overview of the AMP database review and benchmarking workflow. The following steps were taken into account to elaborate this review : (i) systematic identification of AMP databases and AMP predictive tools based on a defined criteria; (ii) construction of a complete dataset by integrating ABP sequences from curated databases, UniProt and sequences retrieved from predictive tools, followed by duplicate removal, sequence standardization, psychochemical annotation, and labeling as ABP or non-AMP; (iii) evaluation of predictive tools using a consistent, reproducible protocol and standard performance metrics, including balanced accuracy, and AUC; (iv) Subgroup analyses per database and tool to assess overall ranking, metric-level performance, and sources of bias, while identifying strengths and limitations related to robustness, generalizability, and reproducibility; and (v) synthesis of key findings into recommendations aimed at improving evaluation guidance, benchmarking practices, and the use of a common sequence dataset for benchmarking AMP prediction.

### Overview of antibacterial peptides and AMP databases

#### Antibacterial peptides mechanisms, properties and challenges

The predominant class of ABPs consists of amphipathic, cationic α-helical structures. These peptides exploit electrostatic complementarity to selectively target the anionic components of microbial lipid bilayers, such as lipopolysaccharides (LPS) in Gram-negative bacteria or teichoic acids in Gram-positive bacteria. Upon sequestration at the membrane interface, the peptides undergo a conformational shift, leading to membrane permeabilization and subsequent osmotic lysis, in order to exert their bactericidal or bacteriostatic effects ^54–56^. However, not all ABPs are cationic, around 12% are classified as anionic or neutral peptides ^57^. This interaction with the bacterial membrane can result in its permeabilization through several distinct models, including the barrel-stave, carpet, and toroidal pore mechanisms ^58,59^. In all cases, the disruption of membrane integrity instigates a lethal cascade, encompassing dissipated ion gradients and the leakage of cytoplasmic contents, and ultimately cell death ^60^. Beyond these primary effects, ABPs frequently exert pleiotropic intracellular activities. Upon translocating across the lipid bilayer, these peptides have been observed to interfere with fundamental processes such as DNA replication, RNA transcription, protein synthesis, and enzymatic catalysis ^61,62^. This multi-target mechanism, which simultaneously compromises structural and metabolic components, presents a significantly higher evolutionary barrier to resistance compared to conventional single-target antibiotics ^10^. Beyond their direct antimicrobial actions, some ABPs exert significant immunomodulatory effects that bridge innate and adaptive immunity. A primary example is the human cathelicidin LL-37, which functions as a bifunctional effector by neutralizing pathogens while simultaneously recalibrating host immune signaling ^63^. These peptides orchestrate complex physiological processes, including leukocyte chemotaxis, the modulation of cytokine expression profiles, and the direction of cellular differentiation. Such multifaceted properties could be of particular therapeutic significance in the management of chronic wounds and persistent inflammatory pathologies where the resolution of immune dysregulation is a clinical priority ^64^.

Despite their therapeutic potential, several physiological and economic constraints currently impede the widespread clinical translation of ABPs. A primary concern remains systemic cytotoxicity, as the biophysical mechanisms targeting bacterial membranes can inadvertently disrupt eukaryotic lipid bilayers, specifically manifesting as hemolysis in erythrocytes ^65–67^. To refine host selectivity, various structural stabilization strategies have been implemented, including peptide cyclization, the incorporation of D-amino acids, and hydrocarbon stapling ^68–70^. Furthermore, the clinical utility of ABPs is often limited by poor proteolytic stability in biological fluids, which severely curtails their systemic half-life and bioavailability ^71,72^. To address these pharmacokinetic deficiencies, advanced delivery platforms such as nanoparticle encapsulation, liposomal carriers, and hydrogel matrices are being deployed to shield peptides from enzymatic degradation and facilitate site-specific release ^73–76^. Finally, the high cost of large-scale solid-phase synthesis remains a significant barrier for complex or elongated sequences ^77,78^. Nonetheless, the advent of optimized recombinant expression systems and innovative bioengineering platforms has begun to address the challenges associated with scalability and production cost ^79–82^.

Despite these hurdles, several ABPs have transitioned into clinical evaluation. However, regulatory approval by the Food and Drug Administration (FDA) and the European Medicines Agency (EMA) remains rare, with only a small number of candidates successfully navigating the clinical pipeline over the past 25 years ^83,84^. A notable success was Daptomycin, a cyclic lipopeptide derived from *Streptomyces roseosporus* that received FDA approval in 2003 for the treatment of complicated skin and skin structure infections. This agent remains a critical therapeutic for combating Gram-positive pathogens, including methicillin-resistant *Staphylococcus aureu*s (MRSA) and vancomycin-resistant enterococci (VRE) ^85,86^. More recently, the FDA approved Dalbavancin in 2014, marking another milestone in peptide-based interventions^87^. Current advancements in synthetic biology and peptide engineering are now facilitating the rational design of novel sequences with optimized antibacterial potency, stability, and pharmacokinetic profiles. These developments represent a transformative shift from the isolation of naturally derived molecules toward the creation of precision-engineered, next-generation peptide therapeutics ^88^.

#### Overview of AMP databases

The primary sequences and physicochemical attributes of AMPs are curated within specialized repositories that facilitate the extraction of specific subsets based on biological origin or functional annotations. However, the inherent structural and functional heterogeneity across the AMP superfamily often hinders accurate classification and impedes direct benchmarking of predictive tools. Recognizing the clinical prominence of ABPs and the need for a common sequence dataset for benchmarking ABPs prediction, we retrieved and integrated all available ABP sequences from the databases summarized in **Table 1** to assemble a comprehensive repository. This unified ensemble incorporates peptides acting against Gram-negative and Gram-positive bacteria to ensure an exhaustive overview of antibacterial activity, providing a more cohesive reference for evaluating current AI-based tools while establishing a refined training foundation for future model development.

**Table 1.**
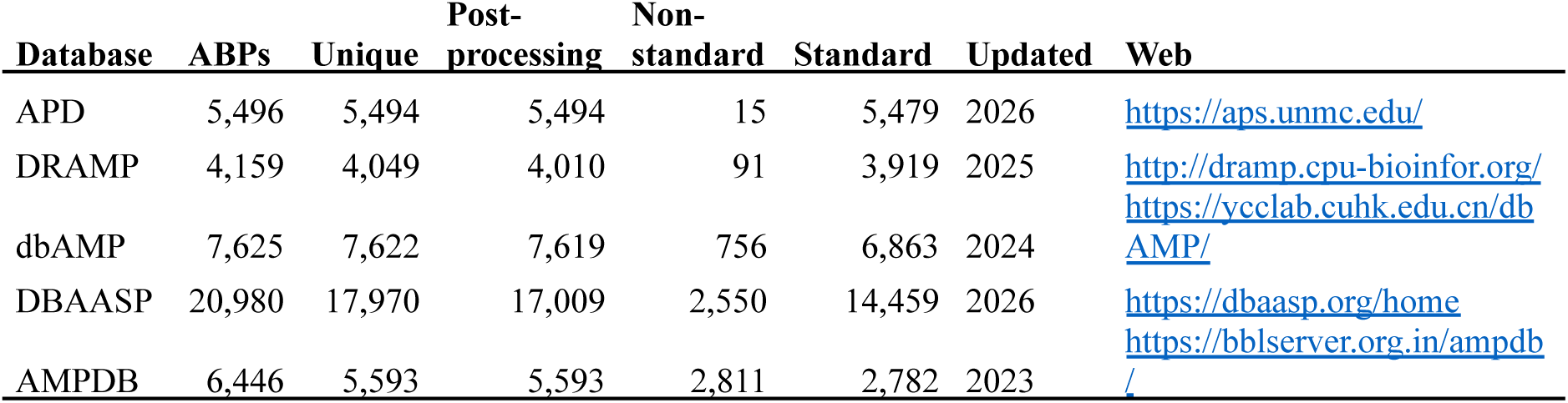
Summary of sequence retrieval metadata across the five selected AMP repositories. Raw sequences underwent standardized preprocessing, including the removal of leading/trailing whitespace, conversion to uppercase, and the elimination of redundant entries. Non-standard sequences were identified and labeled based on length constraints (outside the 5–255 amino acid range), presence of non-alphabetic characters, or inclusion of non-canonical residues. The last update year reported for each database is included. The table was last updated on April 26, 2026. APD, Antimicrobial Peptide Database; DRAMP, Data Repository of Antimicrobial Peptides; dbAMP, Database of Antimicrobial Peptides; DBAASP, Database of Antimicrobial Activity and Structure of Peptides; AMPDB, Anti-microbial Peptide Database

Consequently, the present analysis focuses exclusively on the ABP subclass, applying the rigorous inclusion criteria described in Supplementary file 1 to select the databases that align with the scope of this review. This documentation enumerates both the repositories subjected to evaluation and those that were ultimately excluded for failing to satisfy the established selection standards. To ensure compatibility with the predictive architectures reviewed herein, which exclusively accept the 20 canonical amino acids represented in uppercase format, all retrieved entries underwent standardized preprocessing. This pipeline involved the removal of leading and trailing whitespace characters, conversion to uppercase characters, and the elimination of redundant sequences. The resulting peptide counts, alongside the frequency of non-standard sequences identified during filtration, are detailed in **Table 1**. Comprehensive documentation of the data acquisition process, including the source code and intermediate datasets, is hosted in GitHub (https://github.com/Maximo-7/amp_review).

Established in 2003, the **APD** has evolved from a natural AMP repository into a versatile database that now includes peptides up to 200 residues in length ^13,89^. The database distinguishes between natural, synthetic, and predicted AMPs, provided they exhibit experimentally confirmed antimicrobial functions ^13^. To accommodate a broader range of bioactive polypeptides, the maximum sequence length was recently extended to 200 amino acids. Entries are categorized by their origin into natural, synthetic, and computationally predicted peptides, all of which possess experimentally verified activity. Functional classification within the APD is diverse, covering antibacterial and antifungal types along with various other therapeutic categories. Additionally, the platform integrates computational tools for multiple sequence alignment and AMP prediction. However, a notable technical constraint is that the search and download interface is restricted to FASTA sequences, thereby omitting the rich annotated metadata available on individual entry pages. Integrating these parameters, including minimal inhibitory concentration (MIC) values, defined as the lowest concentration of an antimicrobial compound required to inhibit visible bacterial growth under in vitro conditions; net charge; and the Boman index, an estimate of a peptide’s potential to bind to other proteins, is critical for the rational design of predictive tools. This limitation highlights the need for more comprehensive data extraction methods.

**DRAMP** serves as a comprehensive, curated open-access repository that categorizes AMPs into four distinct subsets: general, patent, clinical, and specific ^15,90^. The patent-derived data provides deep architectural insights through associated patent identifiers and abstracts, while the clinical category archives the progress of peptides currently in the therapeutic pipeline. Within the specific subset, DRAMP organizes specialized sequences into subgroups such as stapled and stability-enhanced peptides. A significant advantage of the DRAMP architecture is the systematic annotation of hemolytic activity and metabolic stability, which are indispensable for the rational design of clinical anti-infectives ^15^. These datasets, in conjunction with other sequence-specific metadata, are available in tabular formats. Furthermore, the platform offers functional integration with fundamental bioinformatics workflows, including sequence alignment and similarity-based searches.

**dbAMP** is a versatile repository that has supported the development of numerous predictive architectures since its inception in 2018 ^14,91^. A distinguishing feature of dbAMP is its integration of structural biology with sequence data. It incorporates predicted tertiary structures for the majority of entries alongside over 100 docked complexes of AMPs and their respective target proteins. These structural models were generated utilizing ESMFold_V1 (Evolutionary Scale Modeling Fold) to provide high-fidelity insights into peptide-target interactions ^92^. The database architecture permits multidimensional filtering based on validation status, biological function, molecular target, toxicity, and structural annotations. Its functional subsets are highly granular, subdividing antibacterial peptides into specialized categories such as antilisterial and antibiofilm agents, among others. However, the utility of the database for large-scale training is hampered by restrictive download capabilities. Currently, the platform only provides a single file with limited metadata, requiring researchers to manually extract detailed annotations for specific peptide subsets. Beyond data hosting, dbAMP also provides integrated computational modules for predicting peptide half-life, hemolytic activity, and antibacterial efficacy.

**DBAASP** represents a foundational resource for structure-activity relationship (SAR) modeling and the *de novo* design of antimicrobial peptides ^93^. Its curated datasets have been instrumental in the development of numerous computational models and derivative databases since 2014 ^94–105^. Unlike platforms that provide binary activity classifications, DBAASP prioritizes quantitative, experimentally determined susceptibility metrics. This includes exhaustive metadata regarding secondary structure, post-translational modifications, and covalent bond architectures, all contextualized within the specific experimental conditions of the assays. A distinctive technical advantage of DBAASP is its programmatic accessibility. Concise summaries are available for manual download, and the entire database is retrievable via a dedicated REST API ^106^ that enables multidimensional filtering across all annotation fields. Furthermore, the platform integrates modules for physicochemical property calculation and species-specific activity prediction. Notably, DBAASP allows for the comparative ranking of entries based on quantitative potency against defined microbial strains or cell lines, a feature that distinguishes it from most extant AMP databases.

**AMPDB** is a recently established, large-scale repository that consolidates peptide data into 88 functional categories based on reported biological activities ^107^. While it is the largest database in this analysis by total entry count, its ABP subset is comparable in size to those of its peers. The platform serves as a meta-repository, aggregating and refining data from the NCBI Protein and EMBL-EBI databases with additional insights from primary literature. Each peptide record is extensively annotated with family classifications, molecular properties, and specific antimicrobial or enzymatic inhibitory functions, often linked via cross-references to external biological databases. A significant technical advantage of AMPDB is its data accessibility, offering structured tabular downloads for both the entire repository and its specific activity-based subsets. Beyond data hosting, the platform facilitates robust sequence analysis through the integration of multiple alignment tools (BLASTp ^108^, MUSCLE ^109^, Needleman-Wunsch ^110^, and Smith-Waterman ^111^) and physicochemical calculators, making it a versatile resource for the preliminary stages of peptide discovery.

### Comparative analysis of antibacterial sequences in AMP databases

For the purpose of this review, we constructed a comprehensive repository by synthesizing data from three primary sources: (i) ABP entries curated from the five aforementioned databases, (ii) non-antimicrobial peptide (non-AMP) sequences retrieved from UniProt (criteria to select sequences from UniProt is defined in Supplementary file 1), and (iii) the training datasets associated with the predictive architectures evaluated in the subsequent sections. This integrated resource, hereafter designated as the complete dataset, contains exhaustive metadata for each sequence. This includes cross-database identifiers, descriptive nomenclature, and boolean attributes for subset classification, such as the designation of standard sequences or evaluation cohorts. Furthermore, the repository archives intrinsic physicochemical descriptors alongside the ground-truth labels and the class assignments generated by each predictive tool. These features are incorporated sequentially; physicochemical properties and model predictions are computed following the assembly of the primary dataset, with tool performance evaluation derived from the final annotated ensemble. The distribution of physicochemical descriptors is of particular significance for comparative database analysis, as detailed in the following sections.

The subsequent analyses focus exclusively on standard sequences, defined as those composed of the 20 canonical amino acids with lengths restricted to a range of 5 to 255 residues. These parameters were established to align with the input constraints of the various AMP prediction tools, ensuring broad compatibility while minimizing data attrition. Regarding the ABP cohort, we observed varying sequence counts across the repositories with significant degrees of overlap. The distribution of unique sequences contributed by each database, alongside all possible combinations, is illustrated in **Figure 2**.

**Figure 2.**
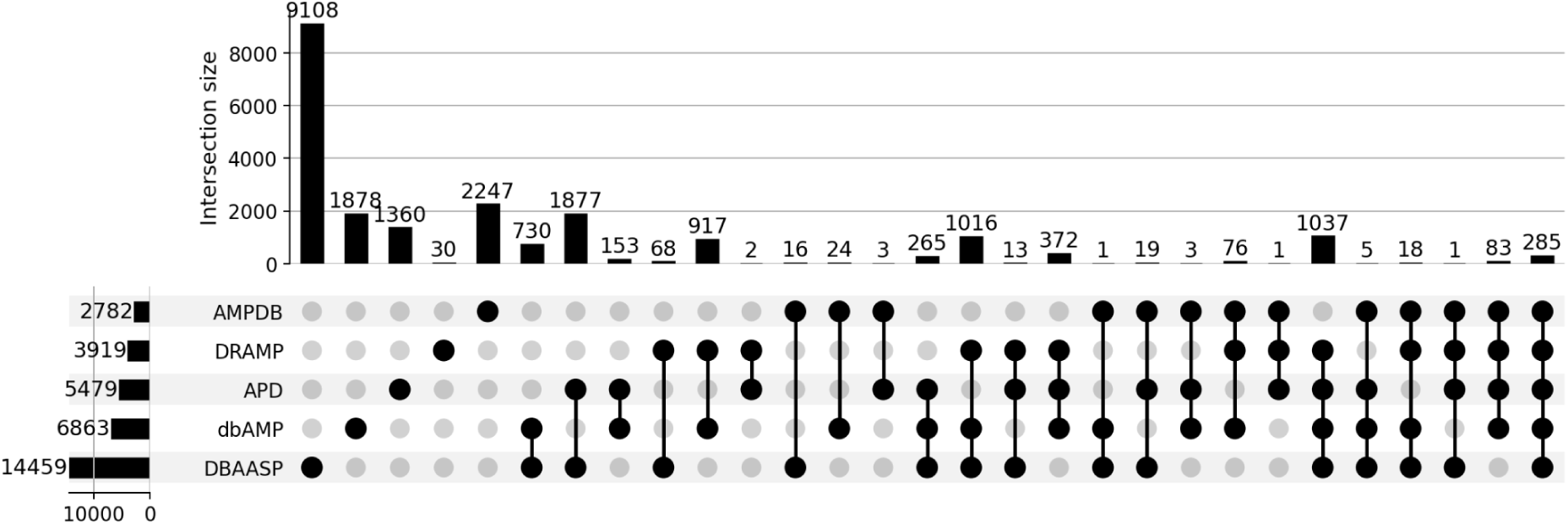
Overlap analysis of ABP sequences across five primary repositories. The UpSet plot illustrates the distribution and overlap of antibacterial peptide sequences within AMPDB, DRAMP, APD, dbAMP, and DBAASP. Vertical bars quantify the cardinality of each intersection across all database combinations, identifying the specific degree of redundancy and uniqueness within the aggregated dataset. Horizontal bars represent the total sequence count provided by each repository, encompassing both unique entries and those shared across multiple platforms. This visualization highlights the consensus and diversity of curated ABP data in the current database landscape.

DBAASP provided the highest absolute volume of unique sequences, with 9,108 entries identified exclusively within its dataset, while DRAMP contributed a marginal set of 30 unique sequences. When normalized by the total number of ABPs in each database, the proportion of unique entries highlights distinct database characteristics; unique sequences accounted for only 0.8% of the DRAMP dataset, compared to 24.8% for APD and 27.4% for dbAMP. Notably, the highest relative uniqueness was observed in DBAASP and AMPDB, where exclusive entries constituted 63.0% and 80.8% of their respective portfolios, suggesting that these repositories harbor a significant amount of primary or non-redundant data not yet integrated into the other major antimicrobial peptide platforms.

A core set of 285 sequences was common to all five databases. Analysis of intersectional cardinality revealed that the largest exclusive pairwise overlap occurred between DBAASP and APD (1,877 sequences). These exclusive intersection values omit sequences that are shared with any additional repositories, ensuring a precise mapping of data redundancy. Significant intersections were also observed among three databases (DBAASP, dbAMP, and DRAMP; 1,016 sequences) and four databases (DBAASP, dbAMP, APD, and DRAMP; 1,037 sequences). While DBAASP acted as a central hub in these intersections, AMPDB remained largely isolated from the dominant shared subsets. To investigate whether these overlaps reflected shared biophysical traits, we performed a comparative analysis of peptide physicochemical properties. The resulting density plots (**Figure 3**) delineate the distributive characteristics of each database. The specific methodologies for calculating these sequence-derived descriptors are documented in Supplementary file 1.

**Figure 3.**
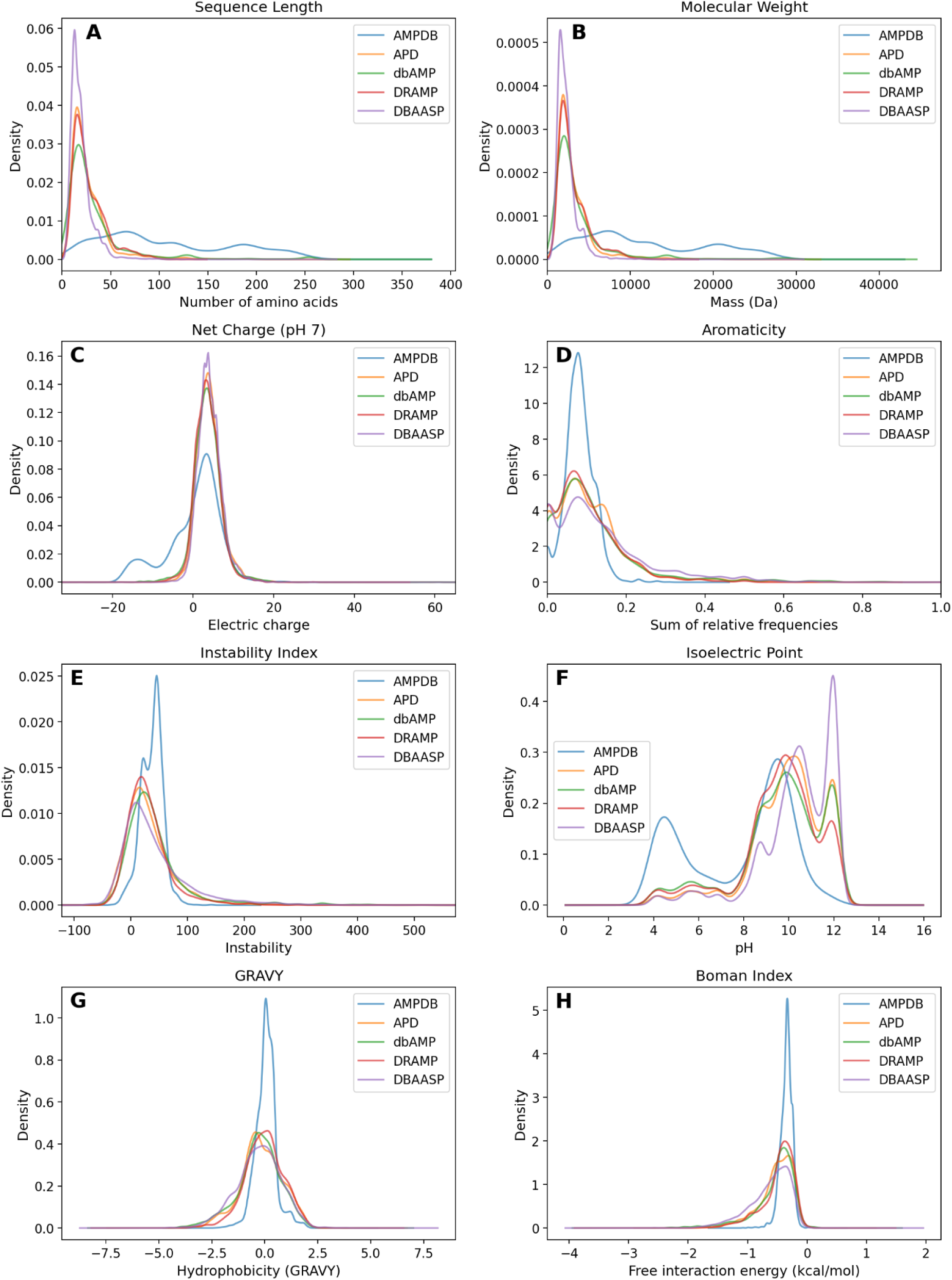
Density distributions of physicochemical attributes across primary ABP repositories. Each panel illustrates the probability density function for a specific physicochemical descriptor derived from standard ABP sequences. The overlaid curves facilitate a comparative assessment of the biophysical landscapes across the five analyzed databases, highlighting regional consensus and divergent property profiles. Variations in these distributions reflect differences in curation priorities and the specialized nature of the curated peptide space within each repository.

The density plots in Figure 3 reveal a consistent distributional overlap among most physicochemical properties across the specialized repositories, with the notable exception of AMPDB, which consistently exhibits divergent patterns. This variance likely stems from its integrative curation model, which draws from general protein databases rather than specific antimicrobial literature. The resulting sequence heterogeneity aligns with the reduced overlaps observed in the Figure 2 intersection analysis. Therefore, AMPDB captures a distinct region of the biophysical landscape, complementing rather than duplicating the other repositories in the aggregated ABP corpus. Sequence length and molecular weight exhibit synchronized distributions across all datasets, confirming the direct linear relationship between these parameters (**Figures 3A and 3B**). Structural constraints within APD (max 200 residues) and DRAMP (max 100 residues) are clearly visible, while DBAASP is uniquely enriched with very short peptides. In contrast, AMPDB explores the sequence space more broadly, showing no significant preference for specific length clusters. A comprehensive analysis of net charge at pH 7 indicates that, in contrast to the prevalence of cationic ABP peptides, the AMPDB database exhibits a notably larger subpopulation of negatively charged peptides compared to other databases (**Figure 3C**). This observation is further corroborated by the appearance of a shoulder on the left side of the isoelectric point plot in AMPDB, which indicates a greater diversity of acidic peptides (**Figure 3F**).

To investigate the distributional anomalies in AMPDB, we performed a secondary analysis of its unique sequences by correlating length with electrostatic profiles. As shown in **Figure 4**, the anionic subset is characterized by a significantly higher mean length (175 residues) compared to the remaining sequences (92 residues). Functional profiling identified a higher prevalence of bactericidal permeability-increasing (BPI) proteins within this long-chain anionic cohort. BPI proteins are a specialized class of membrane-disrupting proteins present in polymorphonuclear leukocyte granules and active against Gram-negative bacteria ^112^. Furthermore, the BPI sequences present in AMPDB exhibited no overlap with entries from the other four databases, none of which included peptides annotated as BPI according to UniProt annotations. The presence of these proteins highlights the potential for significant architectural variation within curated antimicrobial datasets. For sequences with a charge higher than -5 (**Figure 4B**), the length distribution reveals a complex multimodal topology, with a global peak located around 70 residues. The distribution exhibited herein underscores the inherent structural heterogeneity of the dataset, which is evidenced by the presence of diverse peptide classes exhibiting a wide spectrum of chain lengths. Consequently, the characterization of these subpopulations, as illustrated in **Figure 4**, is imperative for the development of next-generation predictive tools. This inherent heterogeneity necessitates a transition toward more granular, feature-aware modeling strategies, which can accurately capture the diverse physicochemical signatures found within expanding AMP families.

**Figure 4.**
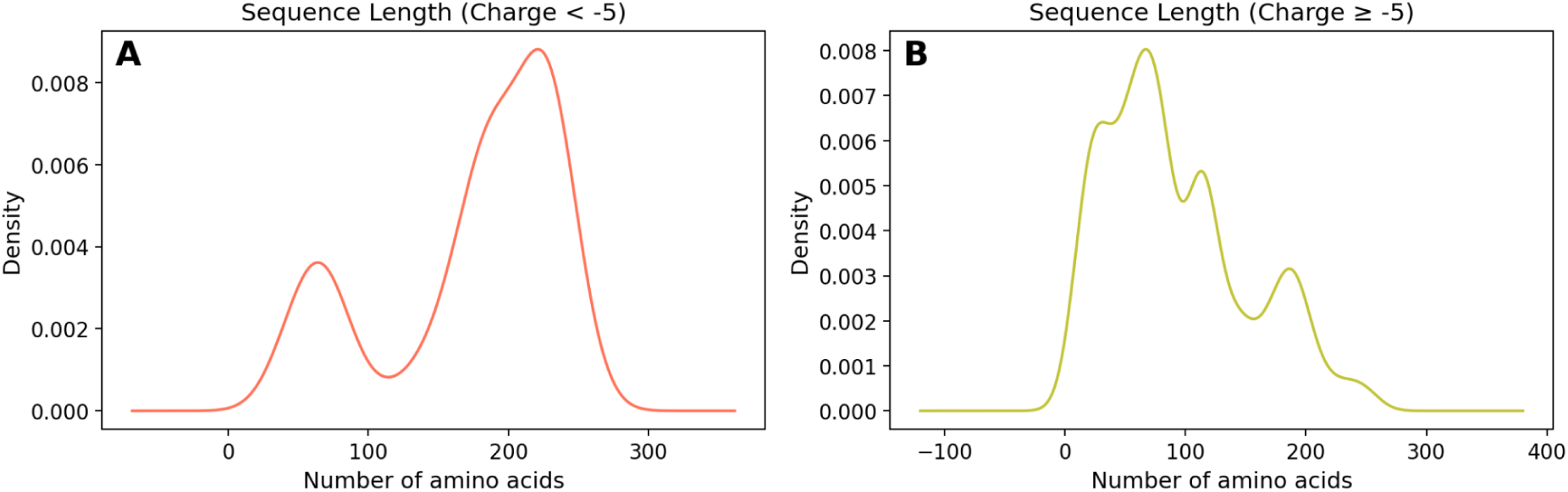
Comparative sequence length density based on a charge threshold from AMPDB. (A) Subpopulation of highly anionic sequences, exhibiting a shift toward higher sequence lengths. (B) Subpopulation of sequences with charge higher than -5, characterized by a multimodal distribution with discrete peaks at approximately 35, 70, 115, 185, and 245 residues.

Analysis of aromaticity and hydrophobicity (GRAVY index) reveals synchronized distributional patterns among the databases, with hydrophobicity curves showing a balanced symmetry around the origin (**Figure 3D and 3G**). In contrast, AMPDB sequences exhibit a significantly lower variance for these properties despite maintaining similar mean values. This constrained dispersion is also reflected in the instability and Boman indices (**Figure 3E and 3H**). Low instability index values across all datasets suggest that the majority of curated ABPs possess moderate predicted structural stability. While the Boman index is typically left-skewed within the [−1.5, 0] range for the majority of databases, except for AMPDB, which exhibits a distinct shift toward more positive values. In addition, the AMPDB dataset exhibits interaction free energy values that lean toward the less negative end of the spectrum, indicating a reduced potential for protein–protein interactions. This observation is consistent with the increased prevalence of negatively charged sequences, which diverge from the canonical cationic AMP profile.

### Comparative analysis of antibacterial and non-antimicrobial peptide sequences

To investigate the distinctive features that separate ABP sequences from non-AMPs, we compared our consolidated ABP dataset against standard non-AMPs extracted from UniProt. The resulting density profiles (**Figure 5**) reveal pronounced distributive shifts in physicochemical properties between the two classes. A two-sample Kolmogorov–Smirnov test was employed to evaluate the null hypothesis of identical distributions. The results confirmed that ABPs and non-AMPs differ significantly across all analyzed properties (p < 10^−283^ in all instances). This comparative framework is critical for characterizing the feature importance and the biophysical decision boundaries utilized by the machine learning architectures evaluated in the following sections. The observed distributional separation suggests that these physicochemical descriptors carry meaningful discriminative signal, contributing to the capacity of classifiers to distinguish ABPs from non-AMPs.

**Figure 5.**
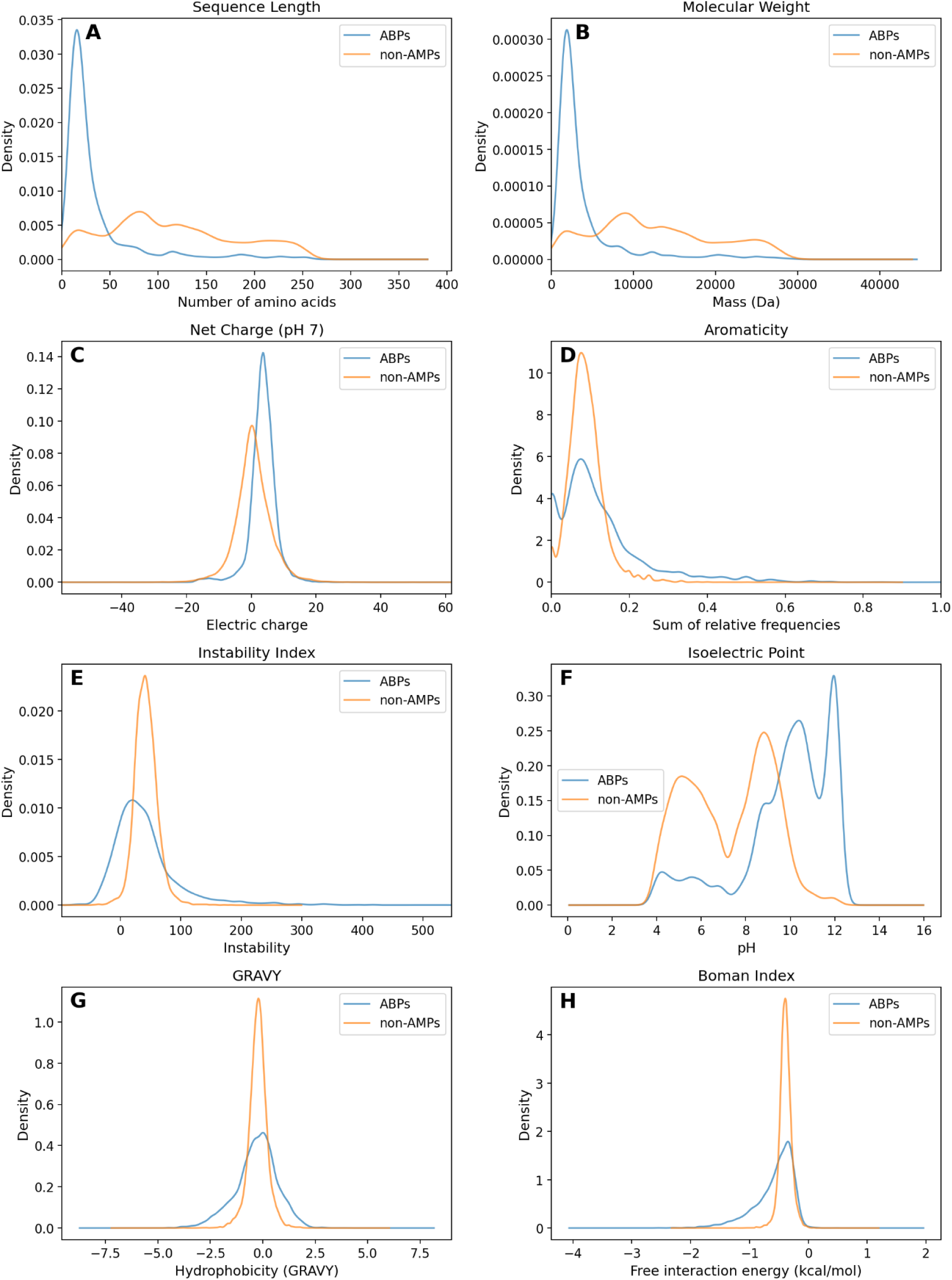
Comparative density distributions of physicochemical attributes for ABPs and non-antimicrobial sequences. Each panel illustrates the probability density function for a specific physicochemical descriptor, comparing the integrated ABP dataset (derived from five primary repositories) against a curated cohort of non-AMPs from UniProt.

Comparative analysis of the sequence space shows that non-AMPs feature higher molecular weight and broader length distributions than ABPs, which consist mainly of peptides under 50 residues (**Figure 5A and 5B**). Although net charge profiles at pH 7 overlap, the non-AMP dataset displays a measurable shift toward anionic values (**Figure 5C**). Isoelectric point data further support this trend because the non-AMP cohort remains biased toward the acidic range despite both sets being bimodal (**Figure 5F**).

Aromaticity values for non-AMPs are concentrated near 0.1, whereas the ABP population is more widely dispersed across the scale (**Figure 5D**). Similar to patterns in the AMPDB repository, non-AMPs show remarkably lower variance in instability, hydrophobicity, and Boman indices than the diverse ABP group (**Figure 5E, 5G and 5H**). While both categories exhibit moderate structural stability, the Boman index reveals a more pronounced negative interaction potential for ABPs. Furthermore, the GRAVY index indicates that ABPs explore a broader and more symmetrical hydrophobicity range centered at zero, a characteristic that supports the formation of amphipathic structures necessary for membrane-active mechanisms

These features collectively suggest that ABPs exhibit a higher propensity for interaction with biological targets and for exerting activity through diverse mechanisms. Their specific cellular targets likely depend on their individual hydrophobic and electrostatic profiles. At the same time, the observed variability across databases suggests the inclusion of functionally diverse peptides with substantially different physicochemical characteristics.

Significant differences between ABPs and non-AMPs are evident across all analyzed properties, pointing to fundamental distinctions rooted in their biological functions. Therefore, these properties serve as highly informative descriptors for class differentiation based on amino acid sequence features. It is important to note that AMPDB sequences show physicochemical distributions more similar to the non-AMP set. This observation indicates the presence of atypical or functionally distinct peptides and underscores the need for rigorous dataset characterization during the development of predictive models.

### Machine learning and deep learning tools for predicting AMPs

The computational landscape for AMP research involves a broad variety of frameworks designed for activity prediction, functional classification, and quantitative estimation of potency (**Figure 6**). These methodologies also extend to the generative design of new sequences. These tools are systematically organized into two main groups according to their modeling techniques. Tools that use classical ML algorithms such as random forest or support vector machine (i) utilize predefined sequence features to capture physicochemical and structural properties. In contrast, modern DL approaches (ii) learn latent representations from raw sequence data or utilize the transformative power of protein language models (PLMs). Many current frameworks adopt hybrid strategies by combining ML and DL components for improved robustness. Additional refinements such as motif scoring and structural constraints are often implemented to guide peptide optimization and ensure biological relevance.

**Figure 6.**
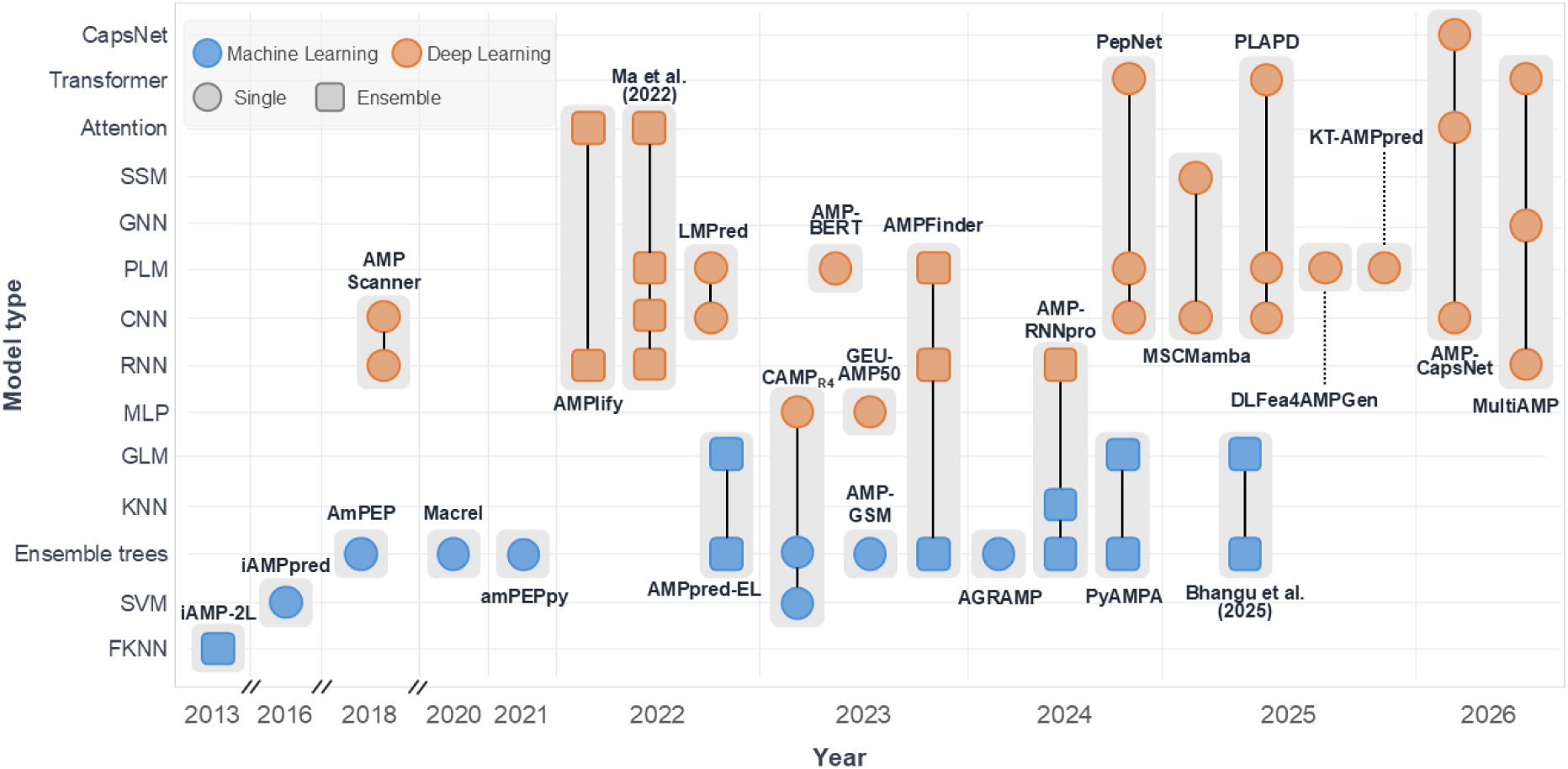
Chronological evolution of computational architectures for AMP prediction. This visualization shows the evolution of predictive models from 2013 to 2026. The Y-axis lists the fundamental neural and algorithmic architectures, while the X-axis indicates the year of publication. The blue and orange markers distinguish between machine learning and deep learning approaches, respectively. Single-model predictors are represented by circles, while squares indicate ensemble methods. Methods that integrate various architectural components are linked by vertical connectors, and the shaded regions delineate the specific components of each tool. The data reveal a significant technological shift from classical machine learning classifiers to advanced deep learning classifiers.

Within the first methodological group (i), early approaches include **iAMP-2L** ^28^ and **iAMPpred** ^29^. iAMP-2L implements a two-layer multilabel classifier based on a fuzzy k-nearest neighbors (FKNN) algorithm: the first layer discriminates AMPs from non-AMPs, whereas the second assigns predicted peptides to one of ten functional categories. In a different approach, iAMPpred employs support vector machines (SVM) combined with information gain-based feature selection to provide specialized classifiers for antibacterial, antiviral, and antifungal activities.

Several subsequent tools rely on tree ensemble models to enhance predictive robustness. **AmPEP** ^30^ uses a random forest (RF) classifier with correlation-based feature selection, while **amPEPpy** ^33^ provides a Python version of AmPEP with a refined model training phase. **AMPpred-EL** ^37^ combines four gradient boosting machine classifiers with a logistic regression meta-model in a two-layer ensemble scheme. **AMP-GSM** ^41^ constructs RF models through a feature grouping and scoring strategy to identify informative descriptor sets for predicting antimicrobial and anti-inflammatory peptides.

Other tools extend this paradigm with additional functionalities. The **CAMP_R4_** ^38^ database offers three separate prediction algorithms, namely a RF, a SVM and a multilayer perceptron, distinguishing between natural and synthetic peptides prior to prediction. **Macrel** ^32^ provides an end-to-end pipeline for AMP discovery from genomic and metagenomic data using a RF classifier; it operates on peptide sequences and raw reads or contigs, and includes a hemolytic activity classifier. Furthermore, **PyAMPA** ^45^ integrates modules for motif identification, RF-based antimicrobial activity against several bacterial species, hemolytic activity and toxicity prediction, half-life estimation via multivariable regression and sequence optimization through mutagenesis and evolutionary algorithms. More specialized tools include **AGRAMP** ^43^, designed to identify peptides active against phytopathogenic bacteria and applied to discover putative AMPs encoded in the citrus (*Citrus sinensis*) genome. More recently, the framework proposed by **Bhangu et al. (2025)** ^48^ combines an extreme gradient boosting classifier with motif ranking and LASSO regression to iteratively generate candidate antimicrobial peptides.

Collectively, the field has undergone a significant evolution from classical machine learning toward sophisticated molecular engineering. This trajectory began with simple sequence discrimination but has progressed into a paradigm of high algorithmic complexity. The adoption of ensemble methods such as Random Forest and XGBoost has proved essential to better manage the inherent noise in biochemical datasets. Furthermore, the systematic integration of toxicity and stability filters has refined the functional granularity of these computational tools, ensuring that safety profiles are evaluated in parallel with antimicrobial potency. Ultimately, the discipline is shifting from passive discovery toward active design. Contemporary frameworks now utilize evolutionary algorithms and regression to optimize the generation of novel peptides for highly specific targets.

Deep learning strategies in the second group (ii) prioritize the extraction of sequence representations directly from raw amino acid data. **AMP Scanner** ^31^ was among the first models to apply deep neural networks to AMP classification, combining convolutional layers with long short-term memory (LSTM) units. A reduced-alphabet variant based on k-means clustering of amino acids was also introduced to simplify the feature space.. Progressively, PLMs pre-trained on large protein databases became central components for AMP classification tasks. **LMPred** ^36^ employed embeddings derived from PLMs pre-trained on large protein databases, including BERT ^113^, T5 ^114^, and XLNet ^115^, each combined with a convolutional neural network classifier. **AMP-BERT** ^39^ uses a fine-tuned BERT architecture for AMP classification and enables interpretation of predictions through self-attention analysis. **PLAPD** ^49^ leverages embeddings from the ESM-2 protein language model and integrates subsequent convolutional and transformer modules for feature extraction. **KT-AMPpred** ^51^ separates AMP identification and functional classification using a fine-tuned T5 PLM and transfer learning for predicting antibacterial, antifungal, and antiviral peptides. In contrast, **GEU-AMP50** ^40^ implements a simple multilayer perceptron with two hidden layers designed to classify short peptides of up to 50 amino acids.

Concurrently, various ensemble and multi-stage architectures emerged. **AMPlify** ^34^ integrates five sub-models incorporating bidirectional LSTM layers with multi-head and context attention modules, and was applied to AMP discovery in the genome of *Rana [Lithobates] catesbeiana*. The pipeline developed by **Ma et al. (2022)** ^35^ combines convolutional networks with LSTM layers and an attention mechanism together with a BERT model to identify novel AMP candidates within human gut microbiome data.. **AMPFinder** ^42^ employs a cascaded architecture in which a RF classifier first identifies AMPs, followed by a deep learning model predicting ten functional classes using PLM embeddings combined with bidirectional LSTM, gated recurrent unit layers and an attention mechanism. **AMP-RNNpro** ^44^ uses stacked ensemble learning in which predictions from multiple machine learning classifiers are used as input to a recurrent neural network meta-classifier. Beyond ensemble strategies, other complex architectures were proposed. **PepNet** ^46^ combines embeddings from a pre-trained T5 model with features extracted via a residual dilated convolution block, feeding a transformer module to predict antimicrobial, anti-inflammatory, and toxic peptides. **AMP-CapsNet** ^52^ introduces a capsule network-based architecture with sequence encoding and channel-attention convolutional modules, using amino acid composition or dipeptide composition as input features.

Finally, some recent works extend beyond functional classification. **MSCMamba** ^47^ focuses on MIC prediction by combining a multiscale convolutional network with a selective state-space model. **DLFea4AMPGen** ^50^ proposes a pipeline for *de novo* peptide generation with potential antibacterial, antifungal and antioxidant properties, based on a BERT-like PLM combined with explainability approaches that are specific from ML. **MultiAMP** ^53^ integrates ESM-2 embeddings with bidirectional LSTM-based sequence modeling and structural features from geometric graph neural networks using cross-attention and transformer modules to jointly predict antimicrobial activity and secondary structure. It also enables rational peptide design and has been applied to identify novel AMPs from ocean-derived proteins. A comprehensive list of the tools assessed, together with associated metadata is provided in Supplementary file 2.

Taken together, these tools illustrate the rapid methodological evolution and diversity of computational approaches for AMP modeling. Although many frameworks report high performance in their modeling tasks, the training corpora used, the variety of AMP activities targeted and the model evaluation protocols vary substantially. Therefore, they are not directly comparable, making it difficult to assess them in a rigorous and fair manner. In addition, functional subtypes within AMPs differ in their physicochemical and biological properties, meaning that predicting each activity effectively constitutes a distinct classification problem. This heterogeneity underscores the importance of systematic benchmarking under standardized conditions and of focusing on a specific AMP class to reduce the intrinsic variability of the underlying datasets. In this review, we therefore concentrate on ABPs, given their clinical relevance and their prevalence among the AMP domain described in the previous sections.

### A standard evaluation framework for AMP predicting tools

For a rigorous and fair evaluation of predictive tools, each method should meet two essential requirements: (i) the datasets used for model development should be publicly available, with training, validation, and test partitions clearly defined; and (ii) the trained prediction model should be directly accessible and usable by independent evaluators. Evaluated tools that failed to meet either of these criteria were flagged accordingly. This approach was strictly followed to mitigate bias and maintain transparency in the interpretation of all evaluation outcomes.

Regarding the first requirement, knowing the exact set of instances used to train each model is essential to construct a rigorous evaluation set that excludes all training examples. Furthermore, the partitioning of the dataset into classes must be clearly specified, ensuring that incompatible classes (e.g., AMPs vs. non-AMPs, or training vs. test sets) do not overlap. Ideally, curated datasets should be deposited in a public repository and linked to the study through a persistent URL or DOI, ensuring long-term availability and consistency with the data described in the publication. Among the 26 tools assessed, only 16 provided their training datasets directly, whereas 13 provided test datasets; and just two, PepNet and MultiAMP, linked their datasets via a DOI. In practice, even nominally accessible data required substantial retrieval effort. The URL reported for AmPEP datasets was inaccessible and the data had to be retrieved through the GitHub repository linked in the Macrel documentation, which reuses those datasets. For PepNet, the linked repository contained several distinct, unclearly labelled corpora, and manual inspection was required to identify which partitions were actually used for training and evaluation. Similarly, the dataset provided by Ma et al. (2022) was not explicitly identified as a training or test partition. In the case of iAMP-2L, the dataset was embedded in PDF supplementary files and had to be extracted manually.

Beyond accessibility, an exhaustive analysis of data leakage, encompassing overlaps between positive and negative classes within a functional category and between training and test partitions, was conducted across all 17 tools for which data were available. Among these, 8 presented overlap between positive and negative classes, with the most affected cases reaching globally overlapping fractions of 22.0%, 3.2% and 1.9%; and 5 presented overlap between training and test partitions, with the largest cases reaching 21.8%, 1.5% and 0.4%. Notably, the tool with the highest class overlap also exhibited the most severe train/test leakage. Complete information on data availability and retrieval for each tool, along with the downloaded datasets, automated download scripts, and data leakage analysis code, is hosted in the project repository. More broadly, the paucity of unique identifiers for peptides underscores the absence of a standardized evaluation framework in this scientific domain, a circumstance that starkly contrasts with the prevailing norm in protein databases.

Regarding the second requirement, the accessibility of predictive models varied considerably across tools, and practical limitations were widespread. Some projects offer web servers that enable rapid prediction, but these often restrict the number of input sequences per submission, as seen with Macrel and AGRAMP, and do not support automation or integration into a standardized benchmarking pipeline. In some cases, such as AGRAMP, results cannot be downloaded and must be manually copied for further analysis. Similar constraints affect tools distributed as local graphical user interface applications, which typically enforce rigid workflows and do not allow isolation of the classification process, as is the case with PyAMPA. Compounding these issues, several web servers referenced in the original publications are no longer accessible, including iAMP-2L, iAMPpred, and AmPEP, while others remain available but with non-functional prediction capabilities, as occurs with AMP-RNNpro. Among the 26 tools evaluated, only 15 provide access to source code, and just nine include trained model files. The absence of trained models is a major obstacle to reproducibility. Even when training protocols and original datasets are available, successful reproduction may be hindered by hardware constraints, software dependency issues, or implementation flaws such as logical errors, missing components, or insufficient documentation.

To address these limitations, we developed individualized Docker environments for each tool, respecting the reported dependency requirements while fixing bugs and standardising the handling of inputs, outputs, and hardware. All preprocessing and prediction steps were integrated into an easy-to-run Nextflow pipeline. Throughout this process, the original logic of each tool was preserved, and published model files were used where available. For the four tools lacking pre-trained models, LMPred, AMP-BERT, PLAPD, and KT-AMPpred, separate training pipelines were also developed using the training parameters specified in the original scripts. Ultimately, of the 15 tools providing source code, only AMP-CapsNet could not be tested: the published code covers only the network architecture definition, omitting all data loading and feature extraction steps as well as any trained model file, making the tool impossible to reproduce or run. The remaining 14 tools were successfully integrated into the Nextflow pipeline. AGRAMP and CAMP_R4_, which could not be incorporated into the pipeline due to the absence of local execution environments, were evaluated separately using their prediction servers. Full details on code adaptations, Docker images, and workflow execution are available in the project repository https://github.com/Maximo-7/amp_review.

A specialized evaluation dataset was extracted from the complete dataset by combining standard ABP sequences from the five primary databases (**Table 1**) with non-AMP entries from UniProt. Potential data leakage was eliminated by the strict exclusion of any instances identified within the training sets of the evaluated tools. The resulting balanced dataset of ABPs and non-AMPs, hereafter referred to as the ‘evaluation dataset’, provided the foundation for our standardized prediction pipeline. This allowed for the extraction of class assignments and probabilities used to annotate the entire dataset. By prioritizing the exclusion of overlapping sequences, this framework establishes a robust and unbiased environment for benchmarking the predictive accuracy of diverse AMP predictive tools.

Analyses of predicted probabilities and classification labels from the evaluation dataset were conducted to assess performance and identify commonalities among the predictive tools (**Figure 7**). This process enabled the identification of significant sequence subpopulations, specifically distinguishing between those that were consistently easy to classify and those that posed persistent challenges for the models. By isolating these subsets, the study delineates the current boundaries of the antimicrobial sequence space that remain difficult to model. Such insights are essential for the targeted improvement of future deep learning and machine learning frameworks. Thus, while some classifiers like DLFea4AMPGen exhibit an AUROC as low as 24.7%, others such as Macrel achieve a more robust 89.9% (**Figure 7A**). These findings suggest that current modeling approaches have yet to reach a level of universal reliability. Furthermore, the observation that many published models perform near random levels during independent testing points to a critical disconnect between self-reported accuracy and rigorous validation. Such results reinforce the importance of adopting transparent and standardized evaluation protocols to accurately assess progress in antimicrobial peptide discovery.

**Figure 7.**
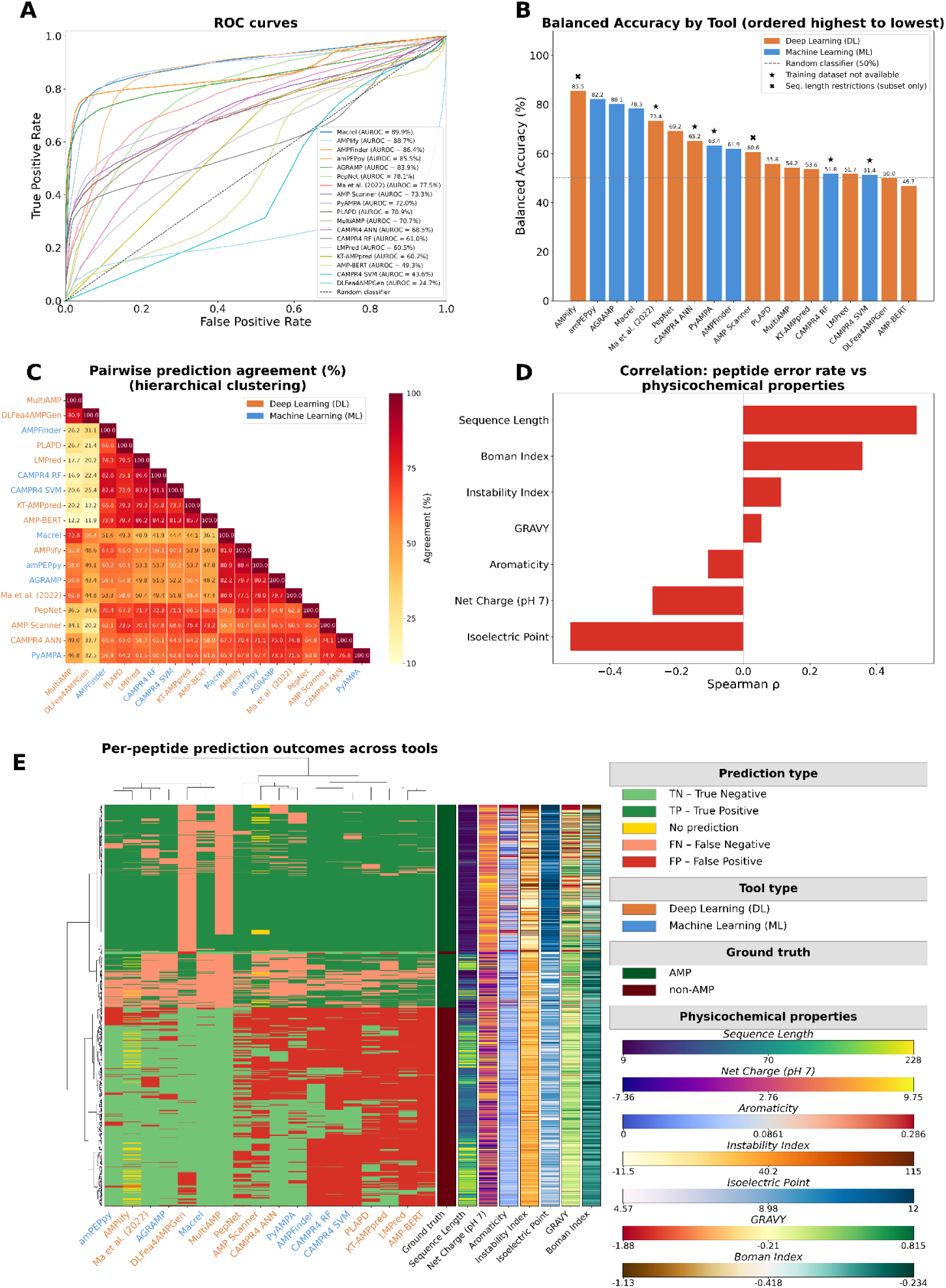
Comprehensive performance and behavioral analysis of 16 antimicrobial peptide (AMP) prediction tools. (A) ROC curves for all 18 evaluated models ranked by AUROC with a dashed diagonal representing the random classifier baseline. (B) Balanced accuracy for each model ordered from highest to lowest. Orange and blue bars distinguish deep learning (DL) and classical machine learning (ML) paradigms respectively. The star symbol (★) denotes tools with non-public training datasets while the cross symbol (✖) identifies tools evaluated on restricted sequence-length subsets. (C) Pairwise prediction agreement matrix organized via hierarchical clustering using average linkage and a distance metric of 100 minus the agreement percentage. Tool labels are color-coded by methodological class. (D) Spearman correlation analysis between the per-peptide error rate and specific physicochemical properties. Red bars signify statistically significant correlations where p < 0.05. (E) Global heatmap of per-peptide prediction outcomes with rows and columns organized by Ward linkage hierarchical clustering. Cell colors indicate true positives (dark green), true negatives (light green), false positives (red), and false negatives (salmon). Yellow cells represent instances where tool-specific length restrictions precluded a prediction. Lateral side strips denote the ground-truth class alongside physicochemical property values clipped to the 5th and 95th percentile range. Specific model configurations were selected for tools offering multiple predictive architectures. For CAMPR4, the ANN, RF, and SVM models were evaluated as separate entities. PyAMPA was represented by the AMPValidate model, while the length/count balanced configuration was utilized for amPEPpy. The LMPred evaluation employed the T5 UniRef50-based model, and AMPFinder was assessed using the stage 1 classifier. For AGRAMP, the 3-gram 9-letter model was selected. The ABP-MPB configuration was used for DLFea4AMPGen, whereas the AMP Fine-tuned Model was chosen for KT-AMPpred. Finally, MultiAMP was evaluated using the sequence-only model. These selections ensure that the most representative or highest-performing available versions of each framework are reflected in the benchmark results.

Figure 7B corroborates this picture while enabling a further comparison by methodological class. In terms of balanced accuracy, although the top-ranked model is DL-based, DL does not emerge as a clearly superior paradigm: the three immediately following positions are occupied by classical ML models, and the ranking as a whole shows no consistent methodological trend. This ambiguity is noteworthy given that DL is the predominant approach in the field, with 11 DL models compared to only seven ML models among those reviewed.

Tools evaluated only on restricted sequence-length subsets occupied ranks 1 and 10 in terms of balanced accuracy, whereas tools for which potential overlap between the training data and the evaluation set could not be explicitly excluded were distributed across ranks 5, 7, 8, 14, and 16. The performance of tools in both categories should be interpreted with caution. For tools with uncertain training data, the evaluation set may include sequences used during model development, introducing a risk of data leakage. For tools assessed on restricted length subsets, the reduced evaluation space may underrepresent difficult-to-classify sequences. In both cases, the reported performance may therefore be inflated relative to that expected under a fully independent and unrestricted benchmark.

Preprocessing methodologies vary significantly across the evaluated frameworks (Supplementary file 2). Deep learning models typically prioritize direct numerical encodings (e.g., one-hot or integer encoding) or PLM embeddings to capture positional and inter-residue relationships via attention mechanisms. In contrast, classical machine learning models frequently employ reduced alphabets based on physicochemical similarity or indirect descriptors to represent the sequence. These distinct strategies suggest that primary linear sequences alone may lack the information density required for high-precision prediction. This potential shortfall highlights either a constraint in current algorithmic feature extraction or a necessity for integrating auxiliary structural and functional annotations. Consequently, future predictive architectures should likely incorporate multi-modal descriptors to improve the classification of complex antimicrobial sequences.

Pairwise agreement across predictive tools is generally low, averaging 59.7% with a peak of 91.1% between the CAMP_R4_ RF and SVM variants (**Figure 7C**). Hierarchical clustering reveals a primary group of high-performing tools, including AMPlify, amPEPpy, AGRAMP, and Macrel, which maintain robust inter-tool agreement above 80%, alongside a significant secondary cluster comprising CAMP_R4_ RF, CAMP_R4_ SVM, KT-AMPpred, and AMP-BERT. In addition, MultiAMP and DLFea4AMPGen form an isolated, highly correlated cluster (80.9%) that nonetheless diverges sharply from all other methodologies. Notably, certain tools demonstrate consistently low agreement with the rest of the cohort. Most strikingly, DLFea4AMPGen records a mean agreement of only 34.7% and appears largely decoupled from the high-performing clusters, showing its highest similarity (56.4%) with Macrel. These findings indicate that while disparate architectures can converge on high accuracy, other models may form clusters due to shared representational strategies or similar training sets. Such patterns suggest that inter-tool agreement is a product of both biological signal and methodological commonalities within the sequence space.

Physicochemical properties significantly influence predictive error rates as quantified by the Spearman correlations in Figure 7D. Sequence length shows the highest positive correlation with failure (ρ ≈ 0.5), while isoelectric point and net charge exhibit the strongest negative correlations. These results demonstrate that models are most accurate when evaluating short, highly basic peptides but fail progressively as sequences become longer or less cationic. Additionally, increased failure rates are associated with higher Boman and instability indices, as well as reduced aromaticity. This systematic bias underscores a foundational limitation in current training methodologies, which prioritize canonical ABP profiles at the expense of more diverse sequences. To enhance the generalization capabilities of future machine learning and deep learning frameworks, training datasets must be expanded to include more uniform distributions across these critical physicochemical parameters, particularly addressing the high error rates associated with non-cationic and high-molecular-weight peptides.

The performance patterns of the evaluated tools are summarized in the global error heatmap shown in Figure 7E, where hierarchical clustering reveals a structured geometric distribution of predictive performance. By dividing the heatmap into six functional sectors based on the intersection of tool profiles and peptide categories, the systematic nature of the limitations of current models becomes evident. The left side of the heatmap is dominated by high-performance tools such as AMPlify, Macrel, amPEPpy, AGRAMP, and the framework by Ma et al. (2022). Within this area, the upper-left sector represents a consensus zone where these models accurately identify canonical ABPs. Below this, in the lower-left corner, a region of maximum specificity is defined. In this sector, the leading tools successfully filter out non-AMPs and demonstrate superior ability to distinguish true negatives even among sequences that mimic cationic properties. However, these same models, despite their overall success, consistently fail on a specific subset of atypical ABPs located in the center-left sector, highlighting a critical scientific frontier. These systematic failures suggest that even state-of-the-art architectures reach an information ceiling when encountering peptides that deviate from classical physicochemical archetypes.

Conversely, the right-hand side encompasses the majority of the evaluated instruments, which are distinguished by substantial performance biases. The upper-right quadrant demonstrates a false consensus, wherein the tools accurately classify numerous ABPs, primarily due to an excessively permissive classification threshold. Notable exceptions to this phenomenon include DLFea4AMPGen and MultiAMP, which exhibit an extremely conservative bias, resulting in the presence of visible vertical bands of false negatives. Moreover, the lower-right quadrant signifies a domain replete with false positives, as over half of the reviewed instruments are incapable of accurately identifying sequences that are not AMPs.

Finally, the center-right sector indicates a blind spot, where tools with a false-positive bias fail to identify the same difficult ABPs as top-tier models. This finding indicates that a non-selective model does not exhibit enhanced biological sensitivity. Rather, it has been demonstrated that the model is capable of recognizing canonical patterns while remaining blind to atypical antimicrobial sequences.

### Conclusions and recommendations

This review has presented a comparative analysis of AMP databases, with a specific focus on ABPs, followed by the application of a standardized framework to evaluate the ability of existing AMP prediction tools to distinguish ABP sequences from non-AMP sequences. Based on our analysis, several conclusions can be drawn, together with specific recommendations to improve both AMP databases and AMP prediction tools.

First, regarding AMP databases, only five out of the 13 resources evaluated met the inclusion criteria of accessibility and active maintenance (Supplementary file 1). This finding highlights the need for sustained funding, curation, and technical support to preserve these resources for the scientific community. Among the five databases retained for analysis, we observed that almost all sequences from DRAMP were also present in other databases. In contrast, the remaining four databases contained a considerable number of unique sequences that were not represented elsewhere. These results indicate that, despite the existence of several AMP repositories, there is still a clear need for a comprehensive ABP-focused database that integrates sequences currently dispersed across different resources.

Furthermore, and perhaps more importantly, AMP databases contain valuable information beyond peptide sequences, including MIC values, structural annotations, toxicity data, target organisms, and experimental validation details. However, this information remains underused for AMP prediction and model development. One major limitation is that most databases primarily allow sequence downloads, while offering limited or no access to structured metadata through APIs, standardized formats, or programmatic query systems. To fully exploit the biological and experimental information contained in these resources, future AMP databases updates should provide interoperable access to curated metadata, including standardized endpoints, downloadable annotation tables, and clear documentation. Such improvements would facilitate the development of more informative prediction tools, enable better benchmarking, and support the design of models that move beyond simple AMP/non-AMP classification toward predictions of antibacterial activity, potency, toxicity, and therapeutic potential.

Several efforts have been made to design and implement new computational tools for AMP detection. However, our analysis reveals a clear discrepancy between the performance reported for many tools in their original publications and their performance under an independent, standardized evaluation framework. This gap underscores the need for external benchmarking based on transparent datasets, consistent evaluation metrics, and independent test sets that are strictly excluded from model training.

Beyond predictive performance, our review also highlights a broader problem of reproducibility and model availability in the field. Only 16 of the 26 tools reviewed were suitable for evaluation, and even within this subset, some tools lacked essential information, such as clearly reported training datasets or ready-to-use models. These limitations make it difficult to determine whether benchmark sequences were previously seen during training, to reproduce the original results, and to compare methods under fair and consistent conditions. Improving the transparency, accessibility, and documentation of AMP prediction tools should therefore be considered a priority for future methodological development.

Therefore, we encourage researchers developing AMP prediction tools to evaluate their models using our benchmark dataset, while ensuring that these sequences, or highly similar homologues, are excluded from their training data. Such practices would reduce the risk of data leakage, improve the reliability of reported performance estimates, and facilitate fairer comparisons among prediction tools.

Taken together, these results lead to several fundamental conclusions. First, there remains considerable room for improvement in the ability of current models to specifically distinguish ABPs from Non-AMPs. This is particularly evident when models are challenged by non-antimicrobial sequences with similar composition or physicochemical properties. This lack of specificity results in the saturation of false positives observed in the right-hand sectors of the global error landscape. Furthermore, the identification of the atypical ABPs located in the center-left sector of the heatmap represents a critical scientific frontier (**Figure 7E**). Because even top-performing models fail systematically in this region, it is clear that current architectures have reached an information ceiling regarding these sequences. Second, further progress in the field will require more carefully designed training corpora that ensure balanced and representative coverage across the full range of relevant peptide properties, especially sequence length, charge, hydrophobicity, and amino acid composition. Such datasets should also minimize redundancy and avoid overlap between training and evaluation sets, as these factors can lead to inflated estimates of predictive performance.

Finally, our results suggest that features derived solely from the linear amino acid sequence may be insufficient to fully capture the biological determinants of antimicrobial activity. Incorporating secondary and tertiary structural information could provide a richer and more biologically meaningful basis for classification. More broadly, these findings indicate that sequence representation may be as critical as the choice of predictive algorithm itself.

In this context, emerging protein language models and structure-aware approaches represent promising directions for future AMP prediction. Their potential impact, however, will depend on the availability of datasets and the ability of researchers to integrate information regarding sequence, structure, and function as a whole.

## Notes

https://github.com/Maximo-7/amp_review/tree/master

https://docs.google.com/document/d/1cR4o7VqmNspxZth382xLRMOGRWJzgN9tmOIC0wyF_a0/edit?usp=sharing

https://docs.google.com/spreadsheets/d/18nAmn41S9xdVHscM_l0m4V-NzIFtORHb/edit?usp=sharing&ouid=115879069630741286767&rtpof=true&sd=true

### Competing Interest Statement

The authors have declared no competing interest.

