## Supplementary file 1 for "Antimicrobial peptide databases and prediction tools: Toward a standard evaluation framework"

#### Criteria to include antimicrobial databases

Criteria: Ability to filter for antibacterial peptides, ongoing updates and maintenance at least every 3 years and accessibility.

##### Databases meeting the selection criteria (Pass)

- APD [1]– last update: January 2026
- DBAASP [2]– last update: 2026
- dbAMP [3]– last update: June 2024
- DRAMP [4]– last update: July 2025
- AMPDB [5]– last update: April 2023
-

##### Databases not meeting the selection criteria (Not Pass)

- CAMP [6]– no updates since 2022; limited accessibility and reliance on predicted data
- PeptideAtlas [7]– inability to filter and download specifically antibacterial peptides
- LAMP [8]– inaccessible; last update in 2013
- ADAM [9]– data download not accessible
- YADAMP [10]– inaccessible , last update uncertain
- ANTIMIC [11]– inaccessible last update uncertain
- SATPdb [12]– data download unavailable; last update uncertain (~2015)
- Bactibase [13]– last update in 2017

#### Methodologies for calculating the sequence-derived descriptors

The physicochemical properties analyzed were inferred directly from the amino acid sequences and are listed below. All properties, with the exception of sequence length and the Boman index, were computed using the *ProteinAnalysis* class from the *Bio.SeqUtils.ProtParam* module of the Biopython library [14].

- Sequence length: the total number of amino acid residues in the sequence.
- Molecular weight: the sum of the molecular masses of all residues in the peptide.
- Net charge (pH 7): the overall electric charge of the peptide at pH 7.
- Aromaticity: a measure of the relative abundance of aromatic residues, defined according to Lobry and Gautier (1994) [15], as the sum of the relative frequencies of phenylalanine (Phe), tyrosine (Tyr), and tryptophan (Trp) in the sequence.

- Instability index: an estimate of protein stability as defined by Guruprasad *et al.* (1990) [16]; values above 40 indicate an unstable protein and are associated with a short half-life.
- Isoelectric point: the pH at which the peptide has a net charge of zero.
- GRAVY (Grand Average of Hydropathy): a measure of the overall hydrophobic or hydrophilic character of the peptide, calculated according to Kyte and Doolittle (1982) [17], as the arithmetic mean of the hydropathy values of all residues in the sequence.
- Boman index: an estimate of the peptide's potential to bind to other proteins, computed as the arithmetic mean of residue-specific interaction free energy contributions along the sequence.

### Criteria to select sequences from UniProt

We used this for searching non-AMP sequences (length:[5 TO 255]) NOT (keyword:KW-0929) NOT (keyword:KW-0211) NOT (keyword:KW-0044) AND (keyword:KW-0964) NOT (keyword:KW-0930) NOT (keyword:KW-0295) NOT (keyword:KW-0878) NOT (keyword:KW-0078) NOT (keyword:KW-0081) NOT (keyword:KW-0425)

Meaning

Do not include these terms: defensin, antimicrobial, antibiotic, antiviral, fungicide, Amphibian defense peptide, Bacteriocin, Bacteriolytic enzyme, Lantibiotic

Include these terms: Secreted, length between 5 and 255
